# *Salmonella* Typhi asparaginase-dependent activation of GCN2 promotes bacterial killing in murine macrophages

**DOI:** 10.64898/2026.03.16.712107

**Authors:** Zachary M. Powers, Michael J. McFadden, Gi Young Lee, Tracey Schultz, Luiza A. Castro Jorge, Daniel F. Edwards, Simon Sanchez-Paiva, Jonathan Z. Sexton, Katherine R. Spindler, Jeongmin Song, Mary X. O’Riordan

## Abstract

Many intracellular pathogens stimulate host cell stress by directly or indirectly causing an imbalance in host nutrients; depletion of amino acid pools in particular can act as a danger signal to infected cells. Using a restrictive host model of *Salmonella enterica* serovar Typhi (*S.* Typhi) infection, we identify early induction the integrated stress response (ISR) by viable bacteria, but not heat-killed bacteria. Genetic deletion of the amino acid sensing ISR kinase GCN2 (also known as EIF2AK4) prevented early ISR activation during *S.* Typhi infection, and murine macrophages lacking GCN2 show impaired bacterial clearance and decreased cytokine output. Supplementation of wildtype C57BL/6 murine macrophages with only the non-essential amino acid asparagine was sufficient to suppress *S.* Typhi-induced ISR activation and deletion of *S.* Typhi *ansB*, encoding an asparaginase, prevented ISR activation during infection. Pharmacological inhibition of mammalian target of rapamycin (mTOR), the other major amino acid sensing pathway in eukaryotic cells, prevented GCN2 activation and ISR induction in murine macrophages, indicating an upstream role for mTOR in signaling to GCN2. These findings suggest a role for the ISR in macrophage innate immune responses to *S.* Typhi infection and highlight a potential difference in nutrient-dependent signaling between the *S.* Typhi-susceptible human host and the restrictive murine host centered around asparagine, mTOR, and GCN2.

## Introduction

Despite an estimated 11-21 million annual cases of typhoidal *Salmonella* worldwide, specific research on *Salmonella enterica* serovar Typhi (*S.* Typhi) remains relatively constrained due to its narrow host range (1). Productive *S.* Typhi infection largely occurs only in the human host, causing systemic typhoid fever, while commonly used murine models do not support infection, leading to development of the closely related *Salmonella enterica* serovar Typhimurium (*S.* Typhimurium) as a more tractable model to study typhoid fever in mice (2, 3). Therefore, murine macrophages can be investigated as a restrictive host model of *S.* Typhi infection to uncover mechanisms that successfully control this pathogen (4, 5). Macrophages are early sentinels of infection, and nutrient availability is central in shaping macrophage innate immune function (6–8). For example, host depletion of tryptophan by indoleamine dioxygenase during *Coxiella burnetti* infection limits bacterial replication (9). Lactate derived from phagocytized bacteria can trigger neutrophil NETosis (10), highlighting how fluctuations in nutrient pools play a key role in alerting the host to infection as an indirect mechanism of pathogen sensing.

Mammalian cells use two primary systems to sense amino acid depletion: mammalian target of rapamycin (mTOR), and the integrated stress response (ISR). Well studied in the context of sterile disease such as cancers, the ISR has a variable role in the immune response to bacterial, viral, and parasitic infections (11–13). Upon activation of any of four ISR stress-sensing kinases (PERK, GCN2, HRI, and PKR), general cellular translation is reduced, through phosphorylation of eIF2α, which allows for preferential translation of *Atf4* mRNA, a transcription factor that generally functions to return the cell to homeostasis or trigger stress-induced cell death (14). Of the four ISR kinases, GCN2 acts as a sensor for amino acid depletion primarily in response to uncharged tRNAs (15, 16). Both Gram-positive and Gram-negative bacterial pathogens induce the ISR during infection (11, 17–20). However, pathogen-induced ISR activation mechanisms and outcomes differ by pathogen, cell type, stimulating signal, and environmental cues, suggesting the specific context of ISR activation influences infection outcomes (21, 22).

Extensive studies establish mTOR as a prominent regulator of cellular metabolism, however, recent findings have revealed new aspects of mTOR signaling and function particularly when associated with lysosomes, a major interface for nutrient acquisition (23–26). mTOR forms distinct complexes, termed mTORC1 (mTOR complex 1) or mTORC2, defined by interacting partners in the complex (27). mTORC1 at the cytosolic-facing lysosomal membrane senses nutrient repletion, promoting growth and inhibiting autophagy (28). mTORC2 activation may occur due to nutrient depletion, repletion, or metabolic waste, with sub-populations localized to the plasma membrane, mitochondria, and endosomes (29, 30). Prior studies demonstrate that mTOR and the ISR can signal bidirectionally in a context-dependent manner, with both pathways capable of regulating transcription, translation, and cellular nutrient pools (31).

As a facultative intracellular pathogen, *Salmonella enterica* activates both intra-and extracellular pathogen-sensing pathways, while perturbing host defenses through effector secretion (32–34). As first responders to early *Salmonella* infection, host macrophages can be co-opted by *Salmonella,* where the bacteria survive inside Salmonella-containing vacuoles (SCVs) and may disseminate to the spleen, liver, and gallbladder through macrophage-dependent and -independent mechanisms (35). Prior studies aimed at identifying human macrophage host factors critical for modulating *Salmonella* infection through a genome-wide CRISPR screen suggested a role for the ISR kinase PKR(36). Together with a prior report that nutrient depletion triggered the ISR during *S.* Typhimurium infection of HeLa cells (11), these findings led us to investigate how host amino acid sensors and the responsive signaling pathways sculpt the macrophage innate immune response during early *S.* Typhi infection of a restrictive host cell.

## RESULTS

### The ISR is induced by live *S.* Typhi during murine macrophage infection

Bacterial infection activates the ISR in a context-dependent way in different host cell types, so we first tested if the ISR was induced during primary murine bone marrow macrophage (BMDMs) infection by *S. enterica* serovars. Macrophages were harvested by whole cell lysis and analyzed by immunoblot for ATF4 protein, which represents the convergence point of the ISR (11, 17–20). Using non-typhoidal *S.* Typhimurium and typhoidal *S.* Typhi infection of BMDMs, we observed that *S.* Typhi but not *S.* Typhimurium infection increased ATF4 levels (**Fig. 1A, B**). Tunicamycin (tun) was used as a positive control for ISR activation as it robustly induces the ISR through endoplasmic reticulum stress via the sensor kinase PERK (21). Treatment of BMDMs with heat-killed bacterial equivalents did not result in increased ATF4 (**Fig. 1A, B**), suggesting an active role for *S.* Typhi in activating the ISR. To orthogonally test this observation, we infected BMDMs with *Salmonella* and measured the number of cells with ATF4-positive nuclei using quantitative immunofluorescence microscopy. Consistent with bulk ATF4 induction observed by immunoblot, we found that only *S.* Typhi-infected samples showed ISR induction (**Figure 1C, D**). These results demonstrate that despite *S.* Typhi and *S.* Typhimurium sharing 89% genetic similarity, these two distinct serovars of *Salmonella enterica* differentially engage the ISR in murine macrophages, allowing us to explore the role that this stress response pathway may play in innate immune responses to *S.* Typhi infection in the context of a restrictive host (37).

**Figure 1:**
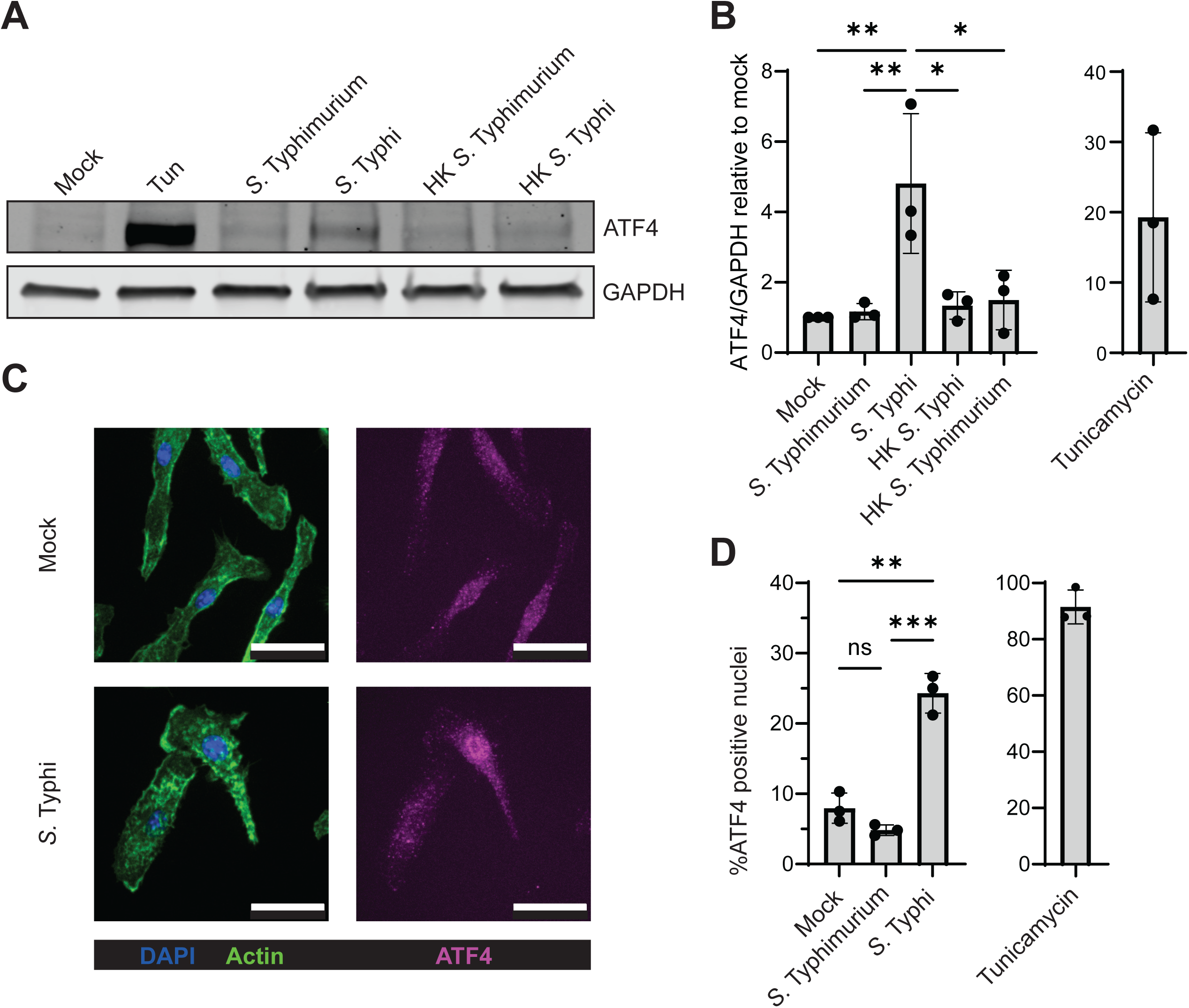
Live *S*. Typhi activates the Integrated Stress Response during murine macrophage infection. WT BMDMs infected with MOI 10 S. Typhimurium or S. Typhi, heat-killed (HK) equivalents, or treated with ISR positive control tunicamycin (tun; 10 µM) for 8h. (A) Whole cell lysates were analyzed by immunoblot for ATF4 as a readout of ISR induction. A representative immunoblot is shown; and 3 experimental replicates were quantified by densitometry and graphed in (B). (C) Representative immunofluorescence microscopy of mock and S. Typhi-infected BMDMs at 8hpi shown as a composite image for DNA (DAPI, blue) and F-actin (phalloidin, green), and a separate image for ATF4 (magenta). Scale bar = 25 µm. (D) Means for ATF4-positive nuclei derived from 3 independent experimental replicates as described in (C). Each point represents the mean of >30 nuclei per replicate sample. Unpaired one-way ANOVA and Tukey9s post-test comparing column means with multiple comparisons. Error bars represent SD. p-value: * < 0.05; ** < 0.01; *** < 0.001.

### *Salmonella* Typhi-driven macrophage ISR activation requires GCN2

We sought to identify the eIF2α kinase responsible for ISR activation during *S.* Typhi infection of BMDMs, first testing the eIF2α kinase GCN2 (also known as EIF2AK4). Guiding us were reports of two closely related Gram-negative bacteria, adherent-invasive *Escherichia coli* and *S.* Typhimurium, triggering GCN2 during *ex vivo* infection of transformed human epithelial cells (11, 19, 38). In mammalian cells, GCN2 is canonically activated by amino acid starvation, primarily sensing increased levels of uncharged tRNAs, and resulting in GCN2 phosphorylation (15, 16). Non-canonical GCN2 activation may result from ribosomal damage or UV-induced ribosomal collisions, although non-canonical activation does not necessarily induce ATF4 (39–41). We tested the necessity of GCN2 for ATF4 induction during *S.* Typhi infection of wildtype (WT) and GCN2-deficient (*Gcn2^-/-^*) BMDMs. Immunoblotting with a phospho-specific antibody that recognizes the activating GCN2 phosphorylation site revealed that *S.* Typhi induced p-GCN2 in infected conditions, and only WT BMDMs exhibited phosphorylated GCN2 upon UV treatment as a positive control **(Fig. 2A, B)**. Additionally, *S.* Typhi-infected *Gcn2*^-/-^ BMDMs did not increase ATF4 levels in response to *S.* Typhi but were still capable of ISR activation when treated with tunicamycin, suggesting that other ISR components remained functional **(Fig. 2A, C)**. We conclude from these observations that GCN2 is the eIF2α kinase required for ISR induction by BMDMs in the early response to *S.* Typhi infection.

**Figure 2:**
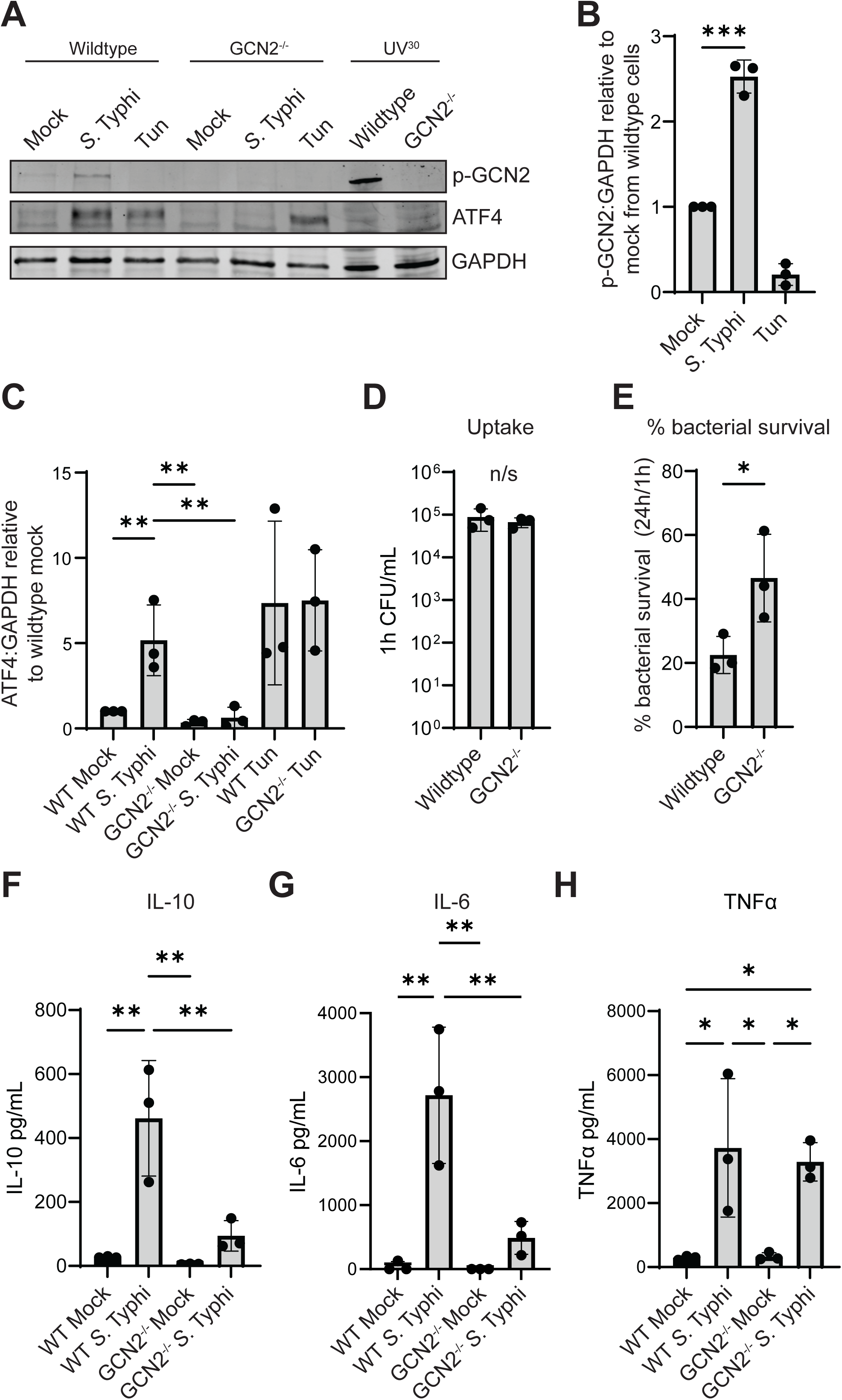
GCN2 is required for *S.* Typhi-driven macrophage ISR activation in murine macrophages. **(A-C)** WT and *Gcn2^-/-^* BMDMs infected with MOI 10 *S.* Typhi for 8hi. Whole cell lysate from ultraviolet (UV) treated BMDM exposed to 500J/m^2^ of energy and recovered for 30 minutes was used as a positive control for phospho-GCN2. **(A)** Representative immunoblot of whole cell lysates subjected to SDS-PAGE and probed with antibodies against phospho-GCN2 and ATF4. **(B)** Densitometry quantification of phospho-GCN2 and **(C)** ATF4 from 3 experimental replicates of (A). Tunicamycin-treated conditions were excluded from analysis. **(D-H)** WT and *Gcn2^-/-^* BMDMs were infected with MOI 10 *S.* Typhi, then lysed and plated for colony forming units to measure **(D)** bacterial uptake at 1hpi and **(E)** 24h bacterial survival (relative to 1hpi). **(F-H)** Supernatants from WT and *Gcn2^-/-^* BMDMs infected with MOI 10 *S.* Typhi were collected at 8h pi and assayed by ELISA for IL-6, IL-10, and TNF³. (B, D, E) Unpaired two-tailed t-test. (C, F-H) Unpaired one-way ANOVA and Tukey9s post-test comparing column means with multiple comparisons. Error bars represent SD. p-value: * < 0.05; ** < 0.01; *** < 0.001.

GCN2 was previously hypothesized to regulate *S.* Typhimurium survival in HeLa cells but its requirement has not been tested directly there or in macrophages (11). Macrophages act early in the innate immune response to phagocytize and kill invading bacteria while signaling to other cells via cytokine secretion. In permissive host-serovar matches, *Salmonella* hijack macrophages and disseminate to distal sites such as the spleen, liver, bone marrow, and gallbladder (42–46). While macrophages generally are not proliferative environments for *Salmonella*, a subset of infected macrophages support bacterial replication, which influences macrophage functional phenotypes (47–49). We tested whether GCN2 was required for *S.* Typhi restriction in WT and *Gcn2^-/-^* BMDMs and found significant differences in the survival of *S.* Typhi after 24h of infection **(Fig. 2D, E)**. We infected *Gcn2^-/-^* and WT C57BL/6 animals with *S.* Typhi, as well as *Gcn2^-/-^Pkr^-/-^*double knockout mice to test for potential redundancy in activating the ISR, but did not observe any significant differences in susceptibility by 24h post-infection when we could still recover *S*. Typhi **(Supplemental Fig. S1A, C, ,F-H)**, potentially indicating more complex interactions *in vivo*. While not statistically significant, levels of circulating cytokines IL-6, IL-10, and TNFα in *Gcn2^-/-^*^-^*Pkr^-/-^*^-^ double knockout mice trended lower during *S.* Typhi infection relative to cytokines from WT mice **(Supplemental Fig. S1I-K)**. To determine if GCN2 was also activated by *S.* Typhi infection in human cells, we infected U-937 human monocyte cells, differentiated with phorbol myristate acetate (PMA). Differentiated U-937 cells did not activate GCN2 during *S.* Typhi or Typhimurium infection, highlighting context-dependent GCN2 activation as a potential contributing factor distinguishing S. Typhi infection in human vs. mouse cells. Tunicamycin treatment increased ATF4, showing that ISR induction remained intact **(Supplemental Fig. S2)**. Treatment of U-937 cells with the GCN2 activator halofuginone did not reduce bacterial burdens after 24h, indicating that GCN2 activation alone is insufficient for bacterial restriction in these human cells **(Supplemental Fig. S3)**. Finally, we returned to murine macrophages to test if GCN2 was required for cytokine production during *S.* Typhi infection. We assayed supernatants from infected BMDMs by ELISA, and found that IL-6 and IL-10, but not TNFα, were reduced in the absence of GCN2 **(Fig. 2F-H)**. Together, these results are consistent with a model where GCN2 enhances the murine macrophage innate immune response to *S.* Typhi by augmenting infection control and potentiating early cytokine production.

### ISR activation can be suppressed by addition of L-asparagine during *S.* Typhi infection

Having established GCN2 as the primary eIF2α kinase driving early ISR induction during *S.* Typhi infection, we next aimed to determine the signal activating GCN2. Since GCN2 is canonically activated in response to uncharged tRNAs in affected cells and infection may alter macrophage amino acid abundance we supplemented amino acids into *S.* Typhi-infected BMDM culture conditions (15, 16, 50). By supplementing in two groups: essential vs. nonessential amino acids, we determined that supplementation with only nonessential amino acids (NEAAs) suppressed *S.* Typhi-induced ISR activation, as measured by ATF4 nuclear intensity using quantitative immunofluorescence microscopy **(Fig. 3A, B, Supplemental Table S1)**. Finding ISR suppression by NEAA supplementation during *S.* Typhi infection was surprising since the previous report of *S.* Typhimurium GCN2 induction in HeLa cells was associated with low levels of the essential amino acids (EAAs) leucine and isoleucine (11). These contrasting findings highlight how the contextual differences between these closely related pathogens and their host environment manifest differently despite both triggering the ISR. To identify the specific amino acid(s) capable of suppressing ISR activation in macrophages, *S.* Typhi and S. Typhimurium BMDM infections were cultured with the addition of individual NEAAs (minus glutamine which was already provided in excess in our standard BMDM culture medium) **(Supplemental Table S2)**, and nuclear ATF4 levels were measured by quantitative immunofluorescence microscopy **(Fig. 3C)**. No *S*. Typhimurium infection conditions showed any significant differences from mock treated. However, of all individual amino acids tested, only the addition of asparagine was sufficient to suppress the ISR in *S*. Typhi-infected macrophages, suggesting that *S.* Typhi-dependent depletion of the NEAA asparagine triggers ISR activation.

**Figure 3:**
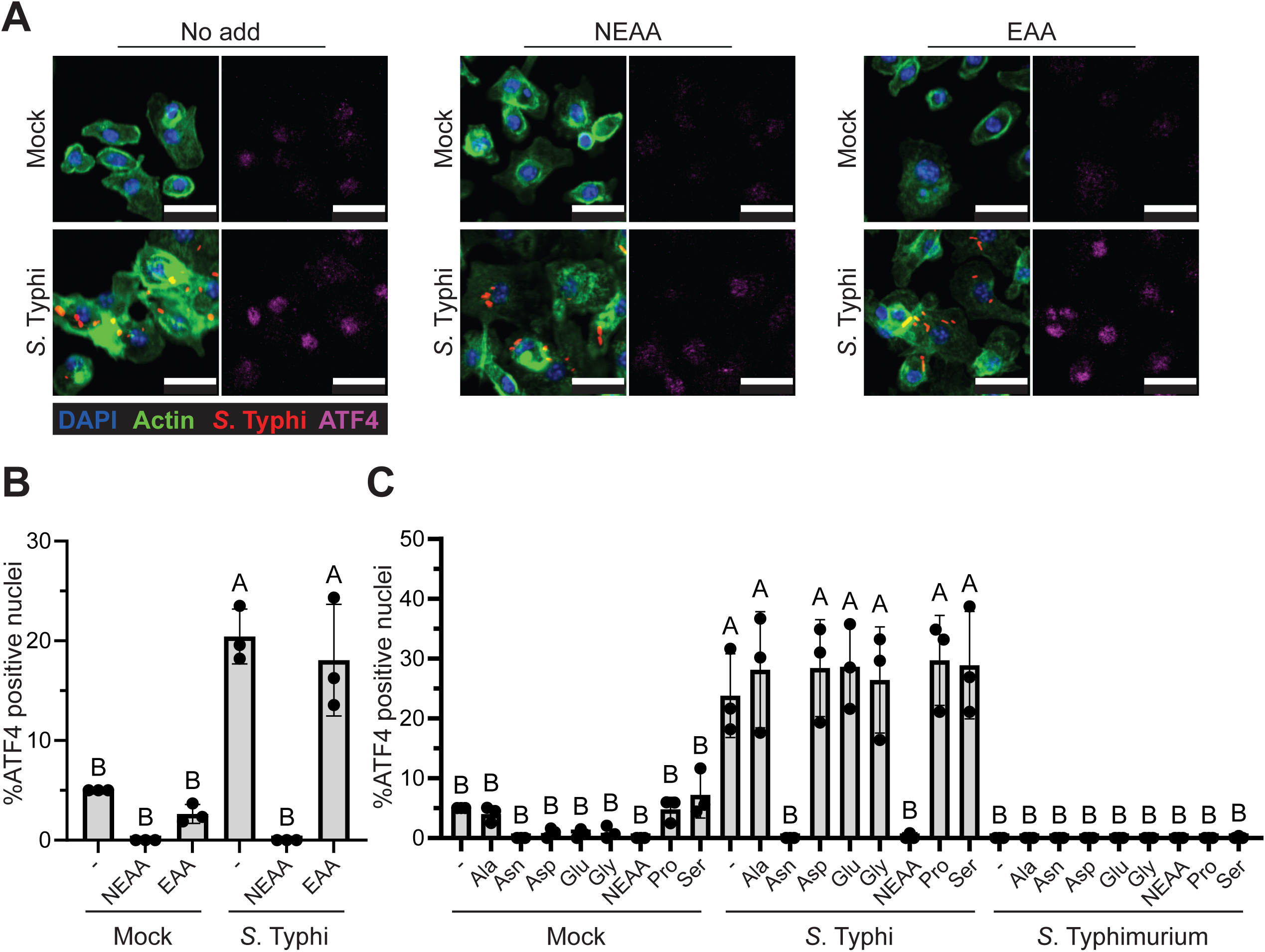
L-asparagine supplementation suppresses *S.* Typhi-induced ISR activation in murine macrophages. **(A-B)** WT BMDMs infected with MOI 10 *S.* Typhi constitutively expressing a DsRed fluorescent reporter were treated with grouped essential- or non-essential amino acids (EAA / NEAA) for 8h. **(A)** Representative immunofluorescence microscopy of ATF4-positive nuclei with composite image including DNA (DAPI, blue), F-actin (phalloidin, green), and *S.* Typhi (DsRed fluorescent protein expression, red), and a separate ATF4 (magenta) image. Scale bar = 25 µm. **(B)** Quantification of 3 experimental replicates of (A) with points representing means of n=3 experiments with >1000 nuclei per condition per replicate sample. **(C)** WT BMDMs infected with MOI 10 *S.* Typhi treated with indicated individual NEAAs during 8h infection. Quantification of ATF4-positive nuclei by immunofluorescence microscopy of S. Typhi-infected BMDMs treated with individual NEAAs. Points represent means of n=3 experiments with >1000 nuclei per condition per replicate. (B, C): Unpaired one-way ANOVA and Tukey9s post-test comparing column means with compact letter display statistical representation of comparisons with p-value < 0.05 (95). For compact letter display, two groups marked with the same letter are not statistically different, whereas two groups marked with different letters, are statistically different. Error bars represent SD.

### The *S.* Typhi L-asparaginase II AnsB is required to induce the ISR during murine macrophage infection

*Salmonella* species encode an asparaginase enzyme and we reasoned that bacterial L-asparaginase (ASNase) might contribute to ISR induction during macrophage infection. Of interest, recombinant ASNases are used as an anticancer treatment to restrict certain types of leukemic growth, and this treatment can activate GCN2 (51). ASNases catalyze the degradation of L-asparagine to aspartic acid and ammonia through hydrolysis and are present in genomes across the tree of life, yet are not expressed in humans or mice outside of the reproductive cells and organs (52–56). *S.* Typhimurium ASNase II, encoded by *ansB*, is expressed and modulates T-cell immune function in a mouse model of infection (57–60). To test if the *S.* Typhi ASNase II is required for ISR induction in murine macrophages, we generated a collection of *S.* Typhi mutants: *ansB* clean deletion (*ΔansB*), *ansB* catalytic-dead point mutant (T111A), *ansB* clean deletion with in-locus complementation with FLAG tag (*ΔansB*+*ansB-F*), and asparagine transporter *ansP* clean deletion (*ΔansP*). Mutants grew similarly to WT *S.* Typhi in axenic culture, and expression in the complemented strain was experimentally confirmed **(Supplemental Fig. S4)**. We performed BMDM infections with these mutants and found that *ansB* deletion or T111A mutation was sufficient to prevent ISR induction during infection, as measured by ATF4 nuclear intensity **(Fig. 4A, B)**. Consistent with Figure 3, supplementation with Asn or NEAAs prevented ISR induction in *ΔansB*+*ansB-F* and *ΔansP* infection conditions **(Supplemental Fig. S5)**. Native locus complementation with *ansB-F* restored ISR induction, though not to WT levels (**Figure 4B)**. These results suggest a model where depletion of asparagine (or an increase in its degradation products within the SCV) induces the ISR. Bacterial mutants were taken up at similar rates to WT *S.* Typhi by WT BMDMs but *ΔansB*+*ansB-F* had slightly higher uptake **(Fig. 4C)**. While we had reasoned that *ΔansB* and *ansB* T111A mutant *S.* Typhi might exhibit increased survival due to reduced ISR activation, instead we found that WT and mutant *S.* Typhi survived similarly at 24h post infection relative to 1h **(Fig. 4C, D)**. We next infected C57BL/6 mice with wildtype and *ΔansB S.* Typhi for 24h to test the requirement for *ansB* during *in vivo* infection and measured bacterial burden in the spleen, liver, and gallbladder 24hpi. *ΔansB S.* Typhi had lower liver burden compared to WT bacteria **(Fig. 4E)** and we observed a trend towards lower burden in the spleen **(Figure 4F)** but did not find statistically significant differences in the gallbladder **(Figure 4G)**, an important reservoir for chronic infection (42). These results suggest *ansB* contributes to *S.* Typhi survival early in infection. Surprisingly, *ΔansB*+*ansB-F* did not fully recapitulate WT *S.* Typhi *in vivo* **(Fig. 4E-G),** despite its capacity to restore ISR activation in macrophages **(Fig. 4A, B)**. Together, these findings establish a role for *S.* Typhi *ansB* in activating the ISR during murine macrophage infection, but also reveal that at least for bacterial killing, deleting bacterial *ansB* was not equivalent to deleting host *Gcn2*. We therefore considered that another amino acid sensor might be engaged during *S.* Typhi infection, with the most likely candidate being the nutrient sensor, mTOR.

**Figure 4:**
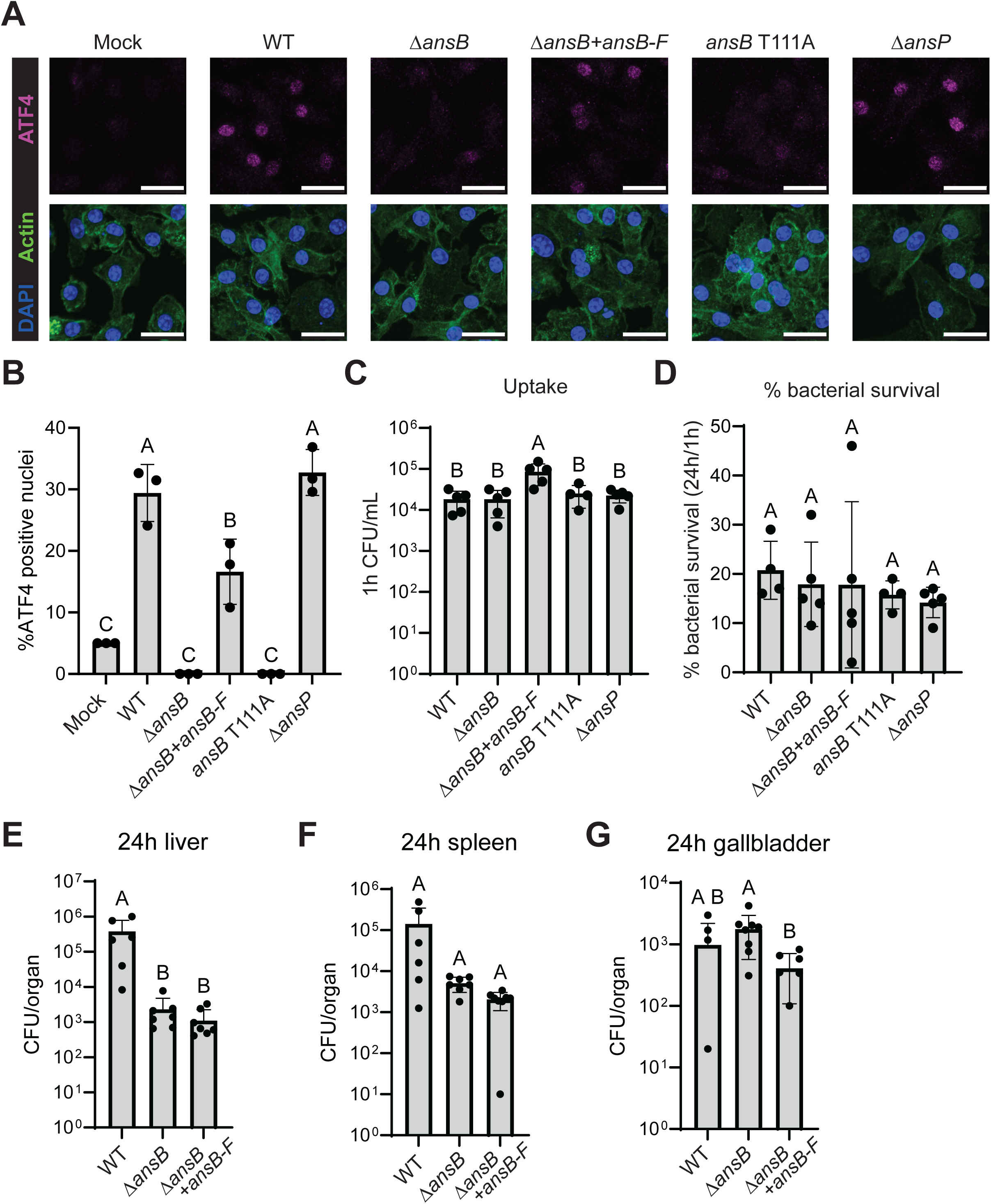
The *S.* Typhi L-asparaginase II AnsB is required for murine macrophage ISR activation during infection. **(A-B)** Murine BMDMs infected with MOI 10 with *S*. Typhi wildtype Ty2, L-asparaginase II (*ansB*) mutants or asparagine transporter (*ansP*) were imaged at 8hpi by automated confocal immunofluorescence microscopy. **(A)** Representative immunofluorescence microscopy of ATF4-positive nuclei with composite DNA (DAPI, blue) and F-actin (phalloidin, green), and with a separate image showing ATF4 (magenta). Scale bar = 25 µm. **(B)** Quantification of 3 experimental replicates of (A) with points representing means of n=3 experiments with >1000 nuclei per condition per replicate. **(C-G)** Murine BMDMs infected with MOI 10 *S.* Typhi wildtype Ty2, *ansB* or *ansP* mutants were lysed and CFU enumerated to measure **(C)** bacterial uptake at 1hpi and **(D)** 24hpi bacterial survival relative to 1hpi. **(E-G)** WT C57BL/6 mice were infected intraperitoneally with 5x10^6^ CFU/animal of *S.* Typhi. At 24h pi, liver **(E)**, spleen, **(F)** and gall bladder **(G)** were harvested, homogenized and CFU enumerated. (B-G): Unpaired one-way ANOVA and Tukey9s post-test comparing column means with compact letter display statistical representation of comparisons with p-value <0.05 (95). For compact letter display, two groups marked with the same letter are not statistically different, whereas two groups marked with different letters, are statistically different. Error bars represent SD.

### mTOR inhibition impairs ISR induction during *S.* Typhi infection

SCVs serve as a niche for *Salmonella* within infected macrophages (61). With 1-3 bacteria within the SCV of most infected cells 8hpi **(Supplemental Fig. S6)**, we questioned whether bacterial asparaginase could deplete cytosolic asparagine sufficiently to induce GCN2, and instead hypothesized that mTOR localized at the SCV membrane might sense asparaginase activity within the SCV. As GCN2 is a known substrate of mTOR, perhaps bacterial ASNase II activity might modulate signaling across the SCV membrane to mTOR to activate cytosolic GCN2(62). To test this hypothesis, we treated *S.* Typhi infected BMDMs with mTOR inhibitors and assayed for p70 S6K phosphorylation (as a proxy for mTOR activation), GCN2 phosphorylation, and ATF4 induction. Treatment with pan-mTOR inhibitor torin 1 completely ablated p70 S6K phosphorylation (Thr389) **(Fig. 5A, B)**, GCN2 phosphorylation, and ATF4 induction **(Fig. 5C-E)**, suggesting mTOR is required in some way to sensitize GCN2 to Asn depletion. Treatment with the mTOR complex 1 (mTORC1) selective inhibitor rapamycin also reduced ISR induction, but not below mock untreated baseline **(Fig. 5C-E)**. Finally, we tested the capacity of *ΔansB S.* Typhi to increase mTOR activity by probing whole cell lysates from infected BMDMs for p-mTOR, p-p70 S6K and p-AKT (Ser473) **(Fig. 5F-L)**. Lack of *ansB* did not alter mTOR phosphorylation compared to WT *S.* Typhi, suggesting AnsB impacts GCN2 through mTOR complex composition or perhaps through an mTOR-independent mechanism **(Fig. 5F-L)**. These results are consistent with a model where mTOR is activated by *S.* Typhi infection of murine macrophages, and is required for GCN2 induction, possibly through spatial regulation of mTOR complexes associated with the SCV. We hypothesize that *S.* Typhi AnsB depletion of asparagine either qualitatively modifies mTOR signaling to promote GCN2 activation or can otherwise activate GCN2 that has been licensed by mTOR **(Fig. 6)**.

**Figure 5:**
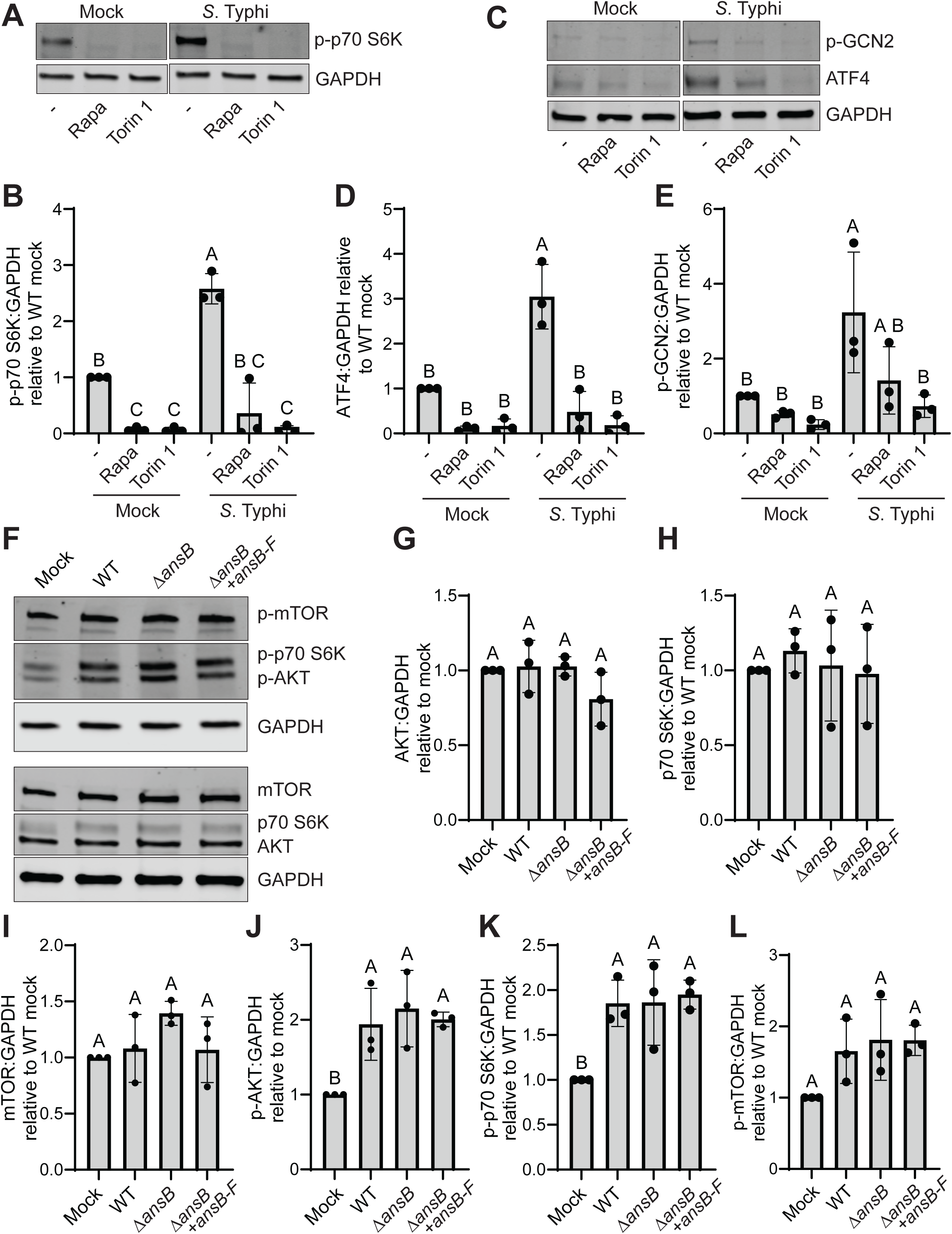
mTOR inhibition prevents *S.* Typhi-induced ISR activation in infected murine macrophages. **(A-E)** WT BMDMs treated with mTOR inhibitors rapamycin (100nM) or torin 1 (250 nM) added at the time of *S.* Typhi infection. Whole cell lysates were harvested at 4hpi followed by SDS-PAGE and immunoblot probed with the indicated antibodies. **(A)** Representative immunoblot of phospho-p70 S6K and **(B)** densitometry quantification of 3 experimental replicates. **(C)** Accompanying representative phospho-GCN2 and ATF4 immunoblots and **(D**, **E)** quantification of 3 experimental replicates of (C). **(F-L)** WT BMDMs infected with *S.* Typhi wildtype Ty2, Δ*ansB* or Δ*ansB+ansB-F* at 4hpi followed by SDS-PAGE and immunoblot probed with the indicated antibodies. **(F)** Representative immunoblot for mTOR, p70 S6K, AKT, and accompanying phosphorylated forms (mTOR Ser2448; p70 S6K Thr389; AKT Ser473). **(G-L)** Densitometry quantification of 3 experimental replicates of (F). B, D, E, G-L: Unpaired one-way ANOVA and Tukey9s post-test comparing column means with compact letter display statistical representation of comparisons with p-value < 0.05 (95). For compact letter display, two groups marked with the same letter are not statistically different, whereas two groups marked with different letters, are statistically different. Error bars represent SD.

**Figure 6:**
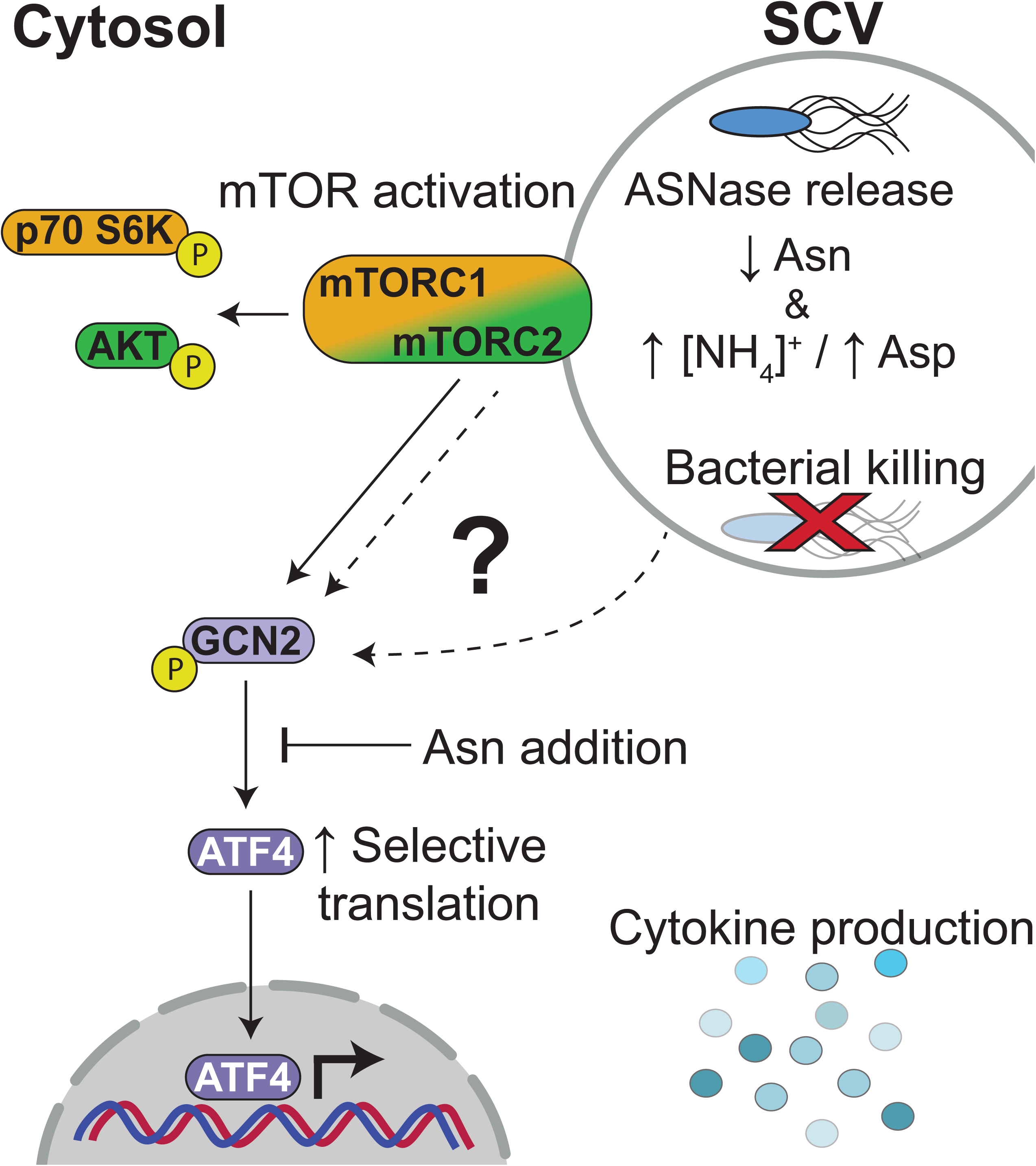
Model of *S.* Typhi ISR activation during murine macrophage infection. We propose the following non-exclusive models for *S.* Typhi induction of the ISR during murine macrophage infection. We hypothesized that the ISR is triggered by bacterial degradation of asparagine in the vacuole, leading to increased aspartic acid (Asp) and ammonia ([NH_4_]^+^) from *S.* Typhi AnsB. In the first model, mTOR at the SCV surface is activated by infection and mTOR signaling is modified in an AnsB-dependent manner, e.g., changing bias from predominantly mTORC1 to mTORC2. The active mTOR complex directly or indirectly phosphorylates GCN2, activating the ISR. GCN2-specific programming promotes cytokine production, and murine macrophages lacking GCN2 have impaired *S.* Typhi restriction. In the second model, infection-induced mTOR licenses GCN2 while *S.* Typhi AnsB acts through an mTOR-independent pathway to induce GCN2 phosphorylation with the same effects on infection outcomes.

## Discussion

Amino acid sensing plays a critical role in cellular homeostasis and the innate immune response to infection. Here, we determine that live *S.* Typhi activates the ISR in a restrictive murine macrophage model of infection via the eIF2α kinase GCN2, dependent on mTOR activation. GCN2 was required to control bacterial infection and potentiate cytokine production early in macrophage infection. U-937 cells did not activate the ISR during *S.* Typhi infection but were capable of inducing the ISR under control conditions, highlighting GCN2 activation as one potential factor that distinguishes the murine vs. human response. We found that GCN2 activation was suppressed or prevented by asparagine supplementation or deletion of the *S.* Typhi ASNase II *ansB* respectively. GCN2-deficient mice remained as resistant to *S.* Typhi infection *in vivo* as wildtype controls. However, deletion of *S.* Typhi *ansB* reduced bacterial burden in the liver of infected animals. *S.* Typhi infection increased mTOR activity as measured by phosphorylation of substrate kinases p70 S6K and AKT, but deletion of *ansB* did not reduce mTOR, p70 S6K, or AKT phosphorylation, suggesting that mTOR may instead play a role in licensing or sensitizing GCN2 to ASNase activity. We therefore favor a model where mTOR activation by *S.* Typhi is tuned by the products of bacterial ASNase activity to induce the ISR to promote macrophage inflammation and host defense **(Fig. 6)**.

Bidirectional signaling between GCN2 and mTOR has previously been documented, but is poorly understood in the context of infection (31, 62–65). We initially proposed a linear hypothesis where bacterial asparaginase activity depleted asparagine, leading to ISR activation. Instead, the results demonstrated a more complex signaling structure with mTOR upstream of ISR induction during *S.* Typhi infection of murine macrophages. Prior studies have established mTOR activation during *S.* Typhimurium infection of human macrophages and epithelial cells and murine macrophages (38, 66–69), including recruitment of mTOR to the SCV through secretion of effectors like SseJ and SopB (70, 71), or activation of the host focal adhesion kinase (67). While *S.* Typhi activates mTOR in human THP-1 cells, this has not previously been reported in murine macrophages (72). Multiple Type III effector proteins secreted by S. Typhimurium are pseudogenes in *S.* Typhi, including *sseJ*, opening up the possibility for serovar-specific mTOR modulation (73, 74). We therefore consider two non-exclusive models where SCV-associated mTOR may directly or indirectly activate GCN2 by sensing AnsB activity through buildup of degradation products aspartate and/or ammonia within the SCV lumen (29, 75). First, *S.* Typhi AnsB activity may determine the functional composition of the mTOR complex, perhaps preferentially activating mTORC2 over mTORC1. Coupled with previous findings that recombinant ASNases impair mTORC1 in some cancer cells(76, 77), we favor a model where mTORC2 signaling at the SCV is primarily responsible for the observed ISR activation during *S.* Typhi infection. Despite evidence of mTORC2 spatially localizing to a subpopulation of vesicles, lack of an mTORC2-specific inhibitor has impeded mechanistic disentanglement of mTORC2 signaling from the ISR and mTORC1 activity (30, 78). Secondly, it is possible that *S.* Typhi infection activates mTOR irrespective of AnsB activity and licenses GCN2 in some way, while AnsB then acts through an mTOR-independent pathway to signal GCN2 activation. It may be that activating GCN2 artificially with halofuginone in human U-937 macrophages did not successfully mimic the coordinated regulation that occurs during infection in the murine macrophages, and therefore did not trigger the anti-microbial function we saw in murine macrophages. Our two proposed models are consistent with observations that during *S.* Typhi infection, both AnsB and mTOR activity are required for phosphorylation of GCN2 and induction of the ISR.

Spatial localization of signaling enables cells to distinguish between extracellular and intracellular environmental cues. Internalization of plasma membrane receptors via phagocytosis allows for combinatorial signaling as machinery of the endolysosomal sensor network, such as mTOR, is recruited to the phagosome. Intriguing work has demonstrated that *S.* Typhimurium ASNase II depletion of L-asparagine during co-culture with T cells is necessary and sufficient to impair T cell function (57, 59, 60). However, T cells do not support *Salmonella* invasion and both WT and *ΔansB* bacteria invade poorly during T cell co-culture (59). In our macrophage model of *S.* Typhi intracellular infection, we find that *S.* Typhi may also shape the innate immune response via ASNase II activity but perhaps using a distinct signaling network. Our data are consistent with a model where a combination of innate immune sensing coupled with the signature of ASNase II activity acts as an alarm for the host cell to identify a state of inflammatory nutrient stress indicative of bacterial infection.

As GCN2 and mTOR both control metabolic reprogramming, it is likely that modulation of these pathways by *S.* Typhi shapes macrophage immunometabolic responses. GCN2 and the ISR exert metabolic control upon activation through transcriptional and translational regulation that cooperate to overcome cellular stress (21). In addition to lipid metabolism and glucose homeostasis, a prominent target of ISR regulation is asparagine synthetase (ASNS) (21, 79). The idea of ASNS regulation by the ISR is intriguing given our finding that *S.* Typhi ASNase II is required for ISR induction, suggesting that perhaps macrophages benefit from sensing depleted asparagine to promote host defense but then must replenish their pools. mTOR also controls many macrophage functions through metabolic and cell cycle regulation (80). Macrophages lacking mTORC1 increase inflammatory cytokine production in response to LPS despite lower glycolytic activity, whereas mTORC2 is required for polarization of alternatively activated M2 macrophages *in vivo* (81, 82). Additionally, when mTOR is inactivated, autophagy is induced to degrade intracellular components such as organelles, damaged proteins, and vesicular cargo to conserve energy and recycle resources (83). Bacterial pathogens like *Salmonella* may be eliminated through autophagic activity and multiple *Salmonella* effectors reinforce mTOR activation to escape autophagy-mediated killing (11, 70, 71). While our data support a GCN2-dependent model for *S.* Typhi restriction downstream of mTOR activation, our model does not rule out bidirectional signaling between GCN2 and mTOR to inactivate SCV-associated mTOR, promoting *Salmonella* through autophagy (63, 64). Future studies will elucidate the complex nutrient sensing mechanisms that coordinate the murine macrophage response to *S.* Typhi infection and determine how engagement of this pathogen with human macrophages may shape the immune environment to be more permissive to chronic infection.

## ACKNOWLEDGMENTS

We thank current and former members of the O’Riordan lab for helpful discussions and support, especially Olivia Harlow and Eliana J. McCray. We also gratefully acknowledge Joel Whitfield and the University of Michigan Rogel Cancer Center Immune Monitoring Core (NIH P30CA046592). This work was supported by the National Institutes of Health, National Institute of Allergy and Infectious Diseases R21AI181115 (M.X.D.O.),F31AI186289 (Z.M.P.), T32AI007528 (Z.M.P., M.J.M), R01AI133935 (K.R.S), the Michigan Pioneer Fellows Program (M.J.M.), the University of Michigan Medical School U081355 (K.R.S), and an S10 grant for high-content imaging equipment S10OD034245 (J.Z.S).

## Methods

### Reagents and consumables

Detailed information for antibodies, reagents and consumables used in this study is found in **Table 1** and **Table 2**.

**Table 1:** Antibodies.

| Target | Catalog number | Manufacturer | Concentration |
| --- | --- | --- | --- |
| ATF4 (D4B8) | 11815 | Cell Signaling | WB: 1:1000; IF: 1:200 |
| p-GCN2 (Thr899) (EPR2320Y) | ab75836 | Abcam | WB: 1:1000 |
| GAPDH (6C5) | sc-32233 | Santa Cruz | WB: 1:5000 |
| FLAG (M2) | F1804 | Sigma-Aldrich | WB: 1:1000 |
| p70 S6K | 9202 | Cell Signaling | WB: 1:1000 |
| p-p70 S6K (Thr389) (108D2) | 9234 | Cell Signaling | WB: 1:1000 |
| AKT (pan) (40D4) | 2920 | Cell Signaling | WB: 1:1000 |
| p-AKT (Ser473) (193H12) | 4058 | Cell Signaling | WB: 1:1000 |
| mTOR (7C10) | 2983 | Cell Signaling | WB: 1:1000 |
| p-mTOR (Ser2448) (D9C2) | 5536 | Cell Signaling | WB: 1:1000 |
| Goat anti-rabbit IRDye 680RD | 926-68071 | LI-COR | WB: 1:5000 |
| Goat anti-mouse IRDye 800CW | 926-32210 | LI-COR | WB: 1:5000 |
| Goat anti-Rabbit IgG (H+L) Cross-Adsorbed Secondary Antibody, Alexa | A21244 | Invitrogen | IF: 1:2000 |
| Fluor™ 647 |  |  |  |

WB = western blot. IF = immunofluorescence.

**Table 2:**
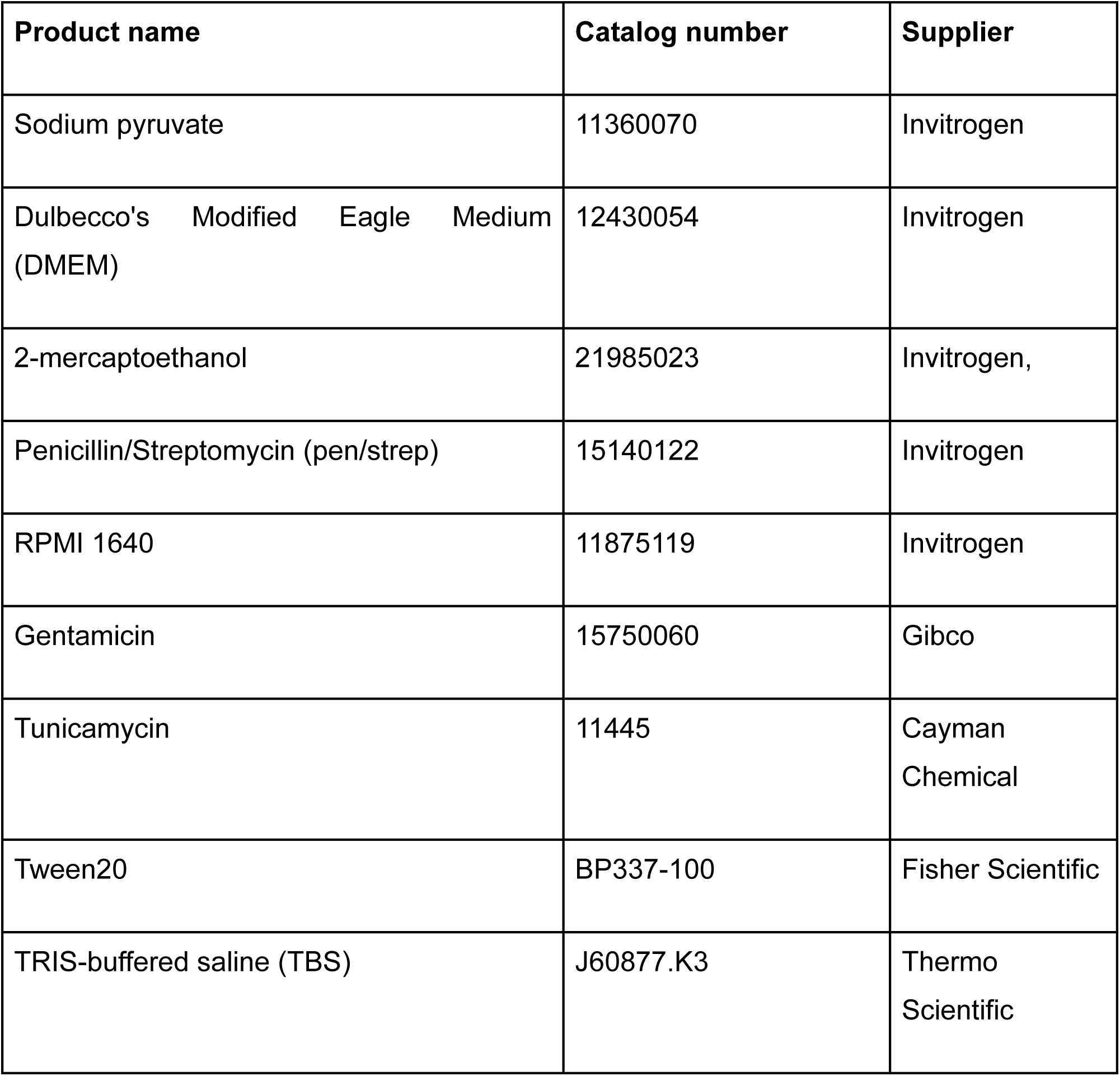
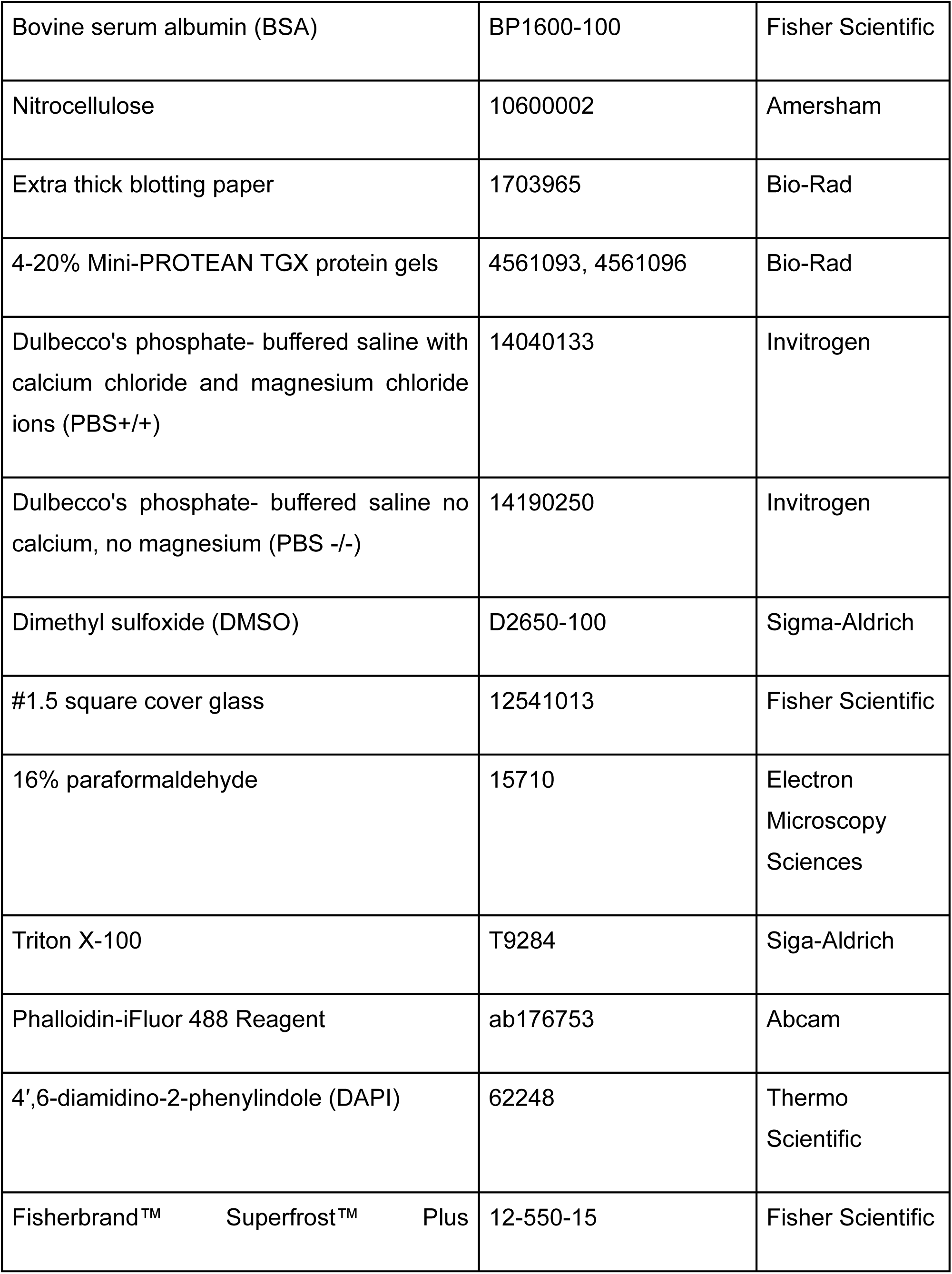
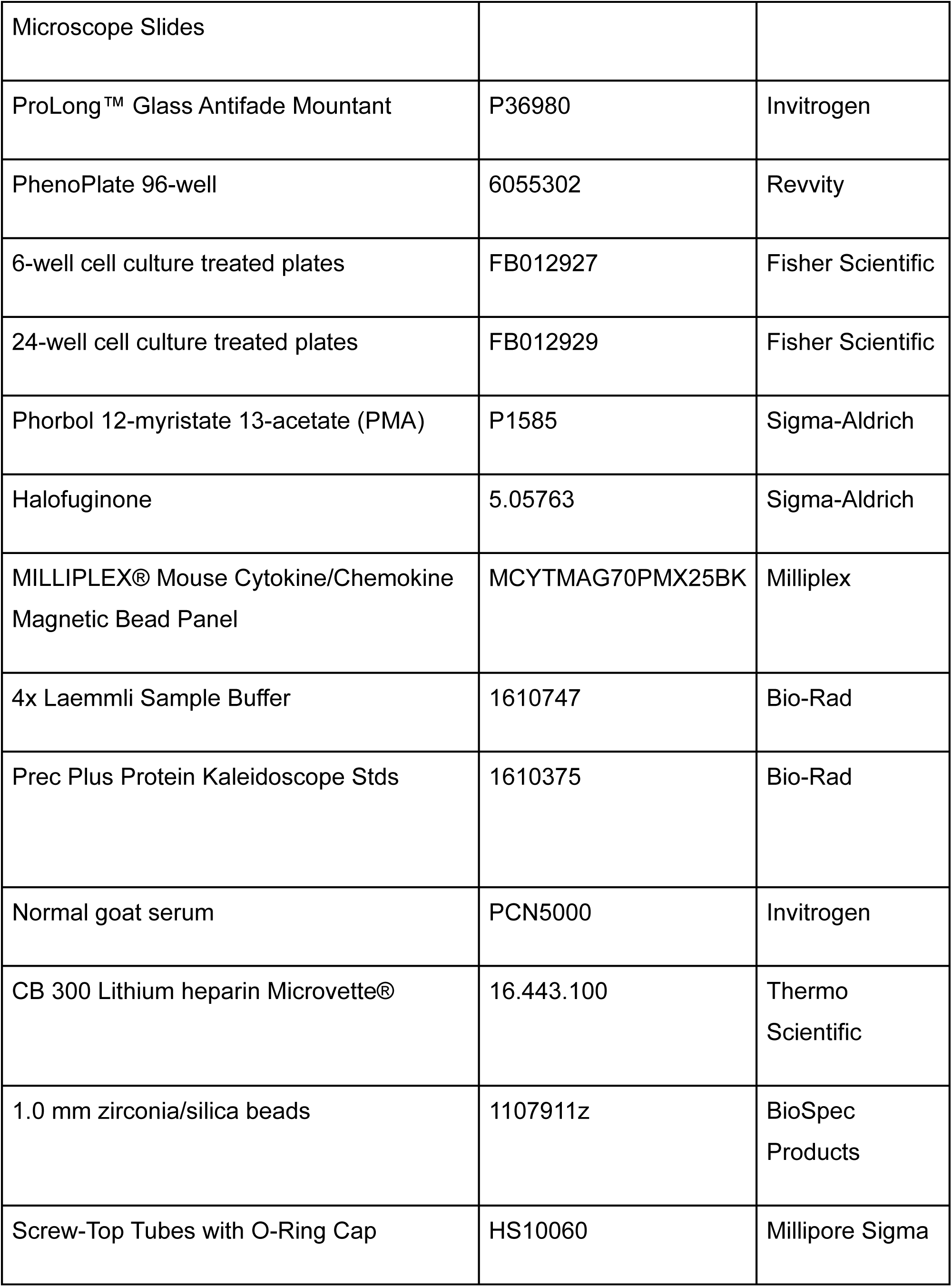
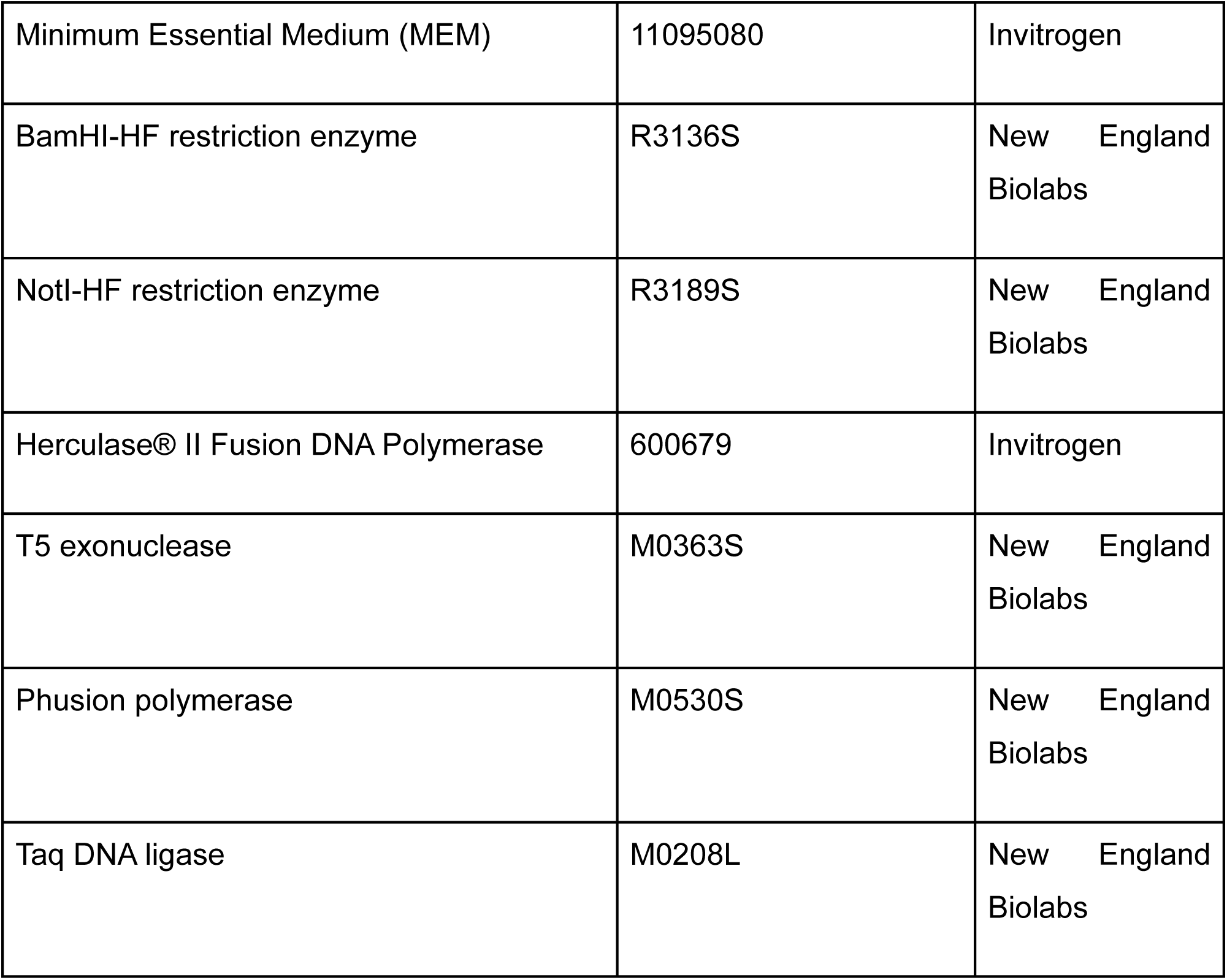
Reagents and consumables.

### Bacterial culture for infections

Bacterial stocks were stored at -80°C in Luria Broth (LB) + 20% glycerol and plated on LB+agar for use with 100 µg/mL ampicillin as needed. A single bacterial colony was inoculated into 3mL of LB with 100 µg/mL ampicillin as needed and grown overnight slanted, shaking, at 37°C. The next day, 1mL of culture was pelleted, supernatant removed and resuspended to known CFU/OD600 in PBS+/+. Each experimental use of bacteria accompanied by CFU plating and overnight incubation to confirm infectious dose. Heat killed bacteria were prepared following PBS resuspension by incubation at 70°C for 1h and killing confirmed by plating on LB+agar and overnight incubation.

### Bacterial cloning

Bacterial mutants were generated as previously described (84–86). Briefly, pSB890 (Tet^R^ and *sacB* for sucrose selection) served as the backbone for gene deletions, point mutations, and FLAG tag insertions. The vector was linearized using restriction enzymes (BamHI-HF and NotI-HF). Inserts for each mutant were amplified by conducting PCR reactions using Herculase® II Fusion DNA Polymerase and specific primers with the *S.* Typhi Ty2 genome as the template. The digested vector and inserts were then assembled using Gibson assembly mix (T5 exonuclease; Phusion polymerase; Taq DNA ligase). The resulting plasmids were transformed into *E. coli* ß2163 Δnic35 for conjugation, followed by subsequent homologous recombination in *S.* Typhi Ty2. All strains were verified by Sanger sequencing through the Cornell Institute Biotechnology Resource Center (BRC) Genomics Facility (RRID: SCR_021727). The locus tag names for *ansB* and *ansP* in the genome of *S.* Typhi Ty2 are T_RS15300 (formerly t3018 or STY3259 in the strain CT18) and T_RS07610 (also known as t1494 or STY1481 in the strain CT18), respectively. The amino acid sequences of AnsB and AnsP can be found at (https://www.uniprot.org/uniprotkb/Q8XGY3/entry and https://www.uniprot.org/uniprotkb/Q8Z739/entry) (84).

### Media formulations

Bone marrow media (BMM): 20% heat inactivated FBS, 30% L-929 conditioned media, 1% sodium pyruvate, 0.1% 2-mercaptoethanol, and remainder DMEM with high glucose, L-glutamine, HEPES, and phenol red. BMM may sometimes be supplemented with 1% penicillin/streptomycin (pen/strep) as noted. L-929 cells were cultured in MEM supplemented with 1% NEAA, 1%, 2 mM L-glutamine, 10 mM HEPES, and 10% FBS. L929-conditioned medium was sterile filtered prior to use.

D10: 10% heat inactivated FBS, 1% Sodium pyruvate, 0.1% 2-mercaptoethanol, remainder DMEM with high glucose, L-glutamine, HEPES, and phenol red.

U-937 complete medium: 10% heat inactivated FBS, 0.1% 2-mercaptoethanol, remainder RPMI 1640.

### Bone marrow isolation and macrophage differentiation

Femurs and tibias from male and female mice aged 8-12 weeks were isolated and bone marrow was extracted by flushing with PBS+/+ with pen/strep. Cells were centrifuged at 250rcf for 5 minutes at 4°C and resuspended in BMM media + pen/strep. Cells were divided into 15cm non-tissue culture treated dishes at a density of ∼3x10^6^ nucleated cells/dish in 25mL BMM + pen/strep. BMM is made with L-929 conditioned media which contains monocytic colony stimulating factor to promote macrophage differentiation (87, 88). For *Gcn2*^-/-^ bone marrow isolation, 1% NEAA was added to BMM (89). Monocytes were differentiated for a total of 6 days, with 30mL of additional BMM + pen/strep added on day 3. On day 6, cells were washed 2x with cold PBS-/-, then 10mL PBS-/- was added and plates were chilled at 4°C for 5 minutes. Cells were then lifted by washing and pelleted at 250rcf for 10 minutes, resuspended in BMM + 10% FBS + 10% DMSO and frozen in 1.5x10^7^ cell/mL aliquots in cryogenic vials. Aliquots were frozen in - 80°C for 24h in a Mr. Frosty freezing container before transferring to liquid nitrogen for long term storage.

### Bone marrow-derived macrophage cell culture and infections

Cells were thawed in a 37°C water bath and diluted 1:10 in prewarmed D10. Cells were pelleted at 250rcf for 5 minutes, counted, plated at 5x10^5^ cells/mL in D10 media in the appropriate cell culture plate (noted per protocol), and incubated overnight at 37°C, 5% CO_2_, with humidity. The following day, the overnight media was removed and replaced with equal volume appropriate for the vessel. Where used, bacteria or heat killed equivalents delivered at MOI 10. Where used as a control, tunicamycin dosed at 5µM. Where used, NEAAs and EAAs were added at 1x final concentration using commercial stocks **(Supplemental Table 1)**. Where supplemented as individual amino acids, stocks were made to match the concentrations of the commercial products found in Supplemental Table 1 and used at 1x final concentration. Following treatment addition, infection was synchronized by centrifugation at 500rcf for 5 minutes. After 30 minutes of incubation, cells were washed twice by PBS+/+, and D10 with gentamicin (50µg/mL) was added. Plates were incubated for an additional 30 minutes, washed twice by PBS+/+, and D10 with gentamicin (5µg/mL) was added. Cells were returned to incubate for the remainder of the experiment as indicated. At time of harvest, cells were washed twice with PBS+/+.

### U-937 cell culture and infections

#### Adaptation of U-937 cells to D10 media

The U-937 cell line used in this study was acquired from ATCC (catalog number CRL-1593.2) circa 2007 and validated using the ATCC Cell Line Authentication Service immediately prior to the submission of this study. U-937 monocytes were acclimated from ATCC recommended complete medium to D10. The U-937 medium recipe contains RPMI 1640 which has nearly 3x the concentration of L-asparagine demonstrated herein to suppress ISR induction (0.378mM vs. 0.100mM, **Supplemental Table 1, 2**), making the switch to D10 necessary and previous studies have demonstrated the capacity for THP-1 cells (another human monocytic cell line) to adapt to DMEM-based media (90). Every second passage, the percentage of D10 to ATCC complete medium was increased by 25%. Doubling time, live cell diameter, and cell viability were measured to ensure cellular health (data not shown). Once 100% D10 was achieved, doubling time recovered to ∼45h within 5 passages and cells were determined to be suitable for use. U-937 cells were maintained between 2.0x10^5^ and 1.0x10^6^ cell/mL and experimental replicates were performed within 5 passages.

#### Infection of U-937 cells

D10-acclimated U-937 cells were plated at 5x10^5^ cells/mL in 2mL D10 media in a 6-well cell culture plate with PMA (100nM) and incubated for 24h at 37°C, 5% CO_2_, with humidity. PMA was removed and cells were washed twice with PBS+/+, then 2mL D10 was replaced and cells were allowed to recover for 24h. The following day, U-937 cells were infected using the same method as BMDMs. Where used, 80nM halofuginone was added for the first two hours of infection before withdrawal by washing twice with PBS+/+ and D10 replacement.

### Fluorescence microscopy

For Figure 1 C-D: Cells were plated in 6-well plates with 5x10^5^ cells/mL in 2mL per well containing glass coverslips and incubated overnight. Following 8h of infection, cells were fixed for 15 mins in freshly prepared 4% paraformaldehyde and permeabilized with PBS+0.1% Triton X-100 (wash buffer). Coverslips were blocked with wash buffer + 3% BSA + 5% normal goat serum (block buffer). Primary ATF4 antibody was incubated overnight at 4°C in block buffer. Coverslips were washed and secondary antibody, phalloidin (1:5000), and DAPI (1:1000) in block buffer were incubated at room temp for 1h in the dark. Coverslips were washed and mounted on glass slides using Prolong™ Glass and cured at room temperature in the dark for 24h before storage at 4°C. Confocal microscopy was performed using Nikon X1 Yokogawa spinning disk microscope at 60x magnification.

All other figures: Fluorescence microscopy was performed as above with the following alterations: cells were plated in 96-well PhenoPlate at 5x10^5^ cells/mL in 0.1mL per well. After washing away secondary staining, wells were stored in 100µL water until imaging. Water was refreshed before imaging on the Yokogawa CQ1 automated high-content spinning disk confocal microscope at 40x magnification. For each image, a 10-slice Z-stack covering the cell height was taken and a max intensity projection was created for analysis with 25-35 fields per well. These conditions resulted in >1,000 cells imaged per condition per experimental replicate.

### Image analysis

Representative microscopy images prepared using FIJI software and using automated macros developed for this study and may be found in Supplemental Software. Briefly, input image pixel intensities are set to user determined values and the representative area of the image is chosen by the user to crop. The cropped selection is saved as single channel .tif and false colored .png files, and a composite false colored .png of non-ATF4 channels (DAPI, phalloidin, DsRed *S.* Typhi). This process is repeated for all representative images in the experiment to keep pixel values across images consistent.

Image analysis was performed with CellProfiler v4.2.8 using analysis pipelines documented here: https://github.com/zmpowers/software-for-GCN2 (91). Briefly, nuclei were identified by DAPI and the cytosol was identified by phalloidin with nuclear masking. Objects touching the image boundary were discarded from analysis. Nuclear ATF4 signal was measured. ROUT analysis was performed on ATF4 nuclear signal with Q=1% and outliers excluded. To calculate % ATF4 positive nuclei, a threshold was calculated using the 95th percentile intensity of mock treated condition per experimental replicate (thereby making 95% of mock treated cell nuclei ATF4 negative). This threshold was propagated to the experimental conditions and the proportion of nuclei with intensities above this threshold are represented as “% ATF4 positive nuclei”. Points represent means of % ATF4 positive nuclei per experimental replicate.

Figure 1D: The nucleus and cytosol were identified as above. Mean ATF4 signal was measured within the nuclei and in the cytosol and a nuclear:cytosolic ratio calculated. Mean nuclear:cytosolic ATF4 signal from the mock treated condition was calculated and used as a threshold. To calculate % ATF4 positive nuclei, this threshold was applied against the nuclear:cytosolic ATF4 signal for each cell and is displayed as a proportion of positive cells analyzed from that condition.

### Bacterial killing and cytokine analysis

Cells were plated in 24-well cell culture plates with 5x10^5^ cells/mL in 0.5mL per well. Cell culture and infection were performed using BMM in place of D10 for killing assays and cytokine collection. At 1h and 24h post infection, supernatants were collected for ELISA analysis and washed twice using PBS+/+before cells lysed in 500µL PBS+1.0% Triton X-100. A 10-fold dilution series was performed, lysates plated on LB+agar, and incubated overnight at 37°C before CFU counting. Enzyme-linked immunosorbent assays (ELISAs) and Luminex cytokine analysis were performed by the UMICH Immune Monitoring Shared Resource core facility.

### Immunoblotting and quantification

Cells were plated in 6-well plates with 5x10^5^ cells/mL in 2mL per well. Following infection, whole cell lysates were collected in 1x Laemmli buffer and stored at -80°C. Before use, samples were heated to 95°C for 5 minutes. Samples were run on 4-20% Mini-PROTEAN TGX protein gels in TRIS-glycine-SDS buffer for 45 minutes at 100V. Proteins were transferred to nitrocellulose membranes using the Bio-Rad Trans-Blot Turbo system (25V limit, 2.5A constant, 20 minutes). Membranes were blocked for 30mins at room temperature in 5% BSA in 1x TBS. Primary antibodies were incubated overnight at 4°C in TBS+5% BSA using antibodies found in **Table 1**. Following incubation, membranes were washed with TBS+0.1% Tween20 (TBS-Tw). Secondary antibodies were added in TBS-Tw+5% BSA and incubated 1h at room temperature. Blots were washed with TBS-Tw before imaging (LI-COR Odyssey Classic, medium quality, 84μm). Quantification was performed using FIJI image analysis software (92). Briefly, the mock lane was selected using the rectangle tool and a box of the same size was propagated to all lanes. Peaks of interest were chosen by drawing a line at the inflection point on either side of the peak base and the area under the curve was measured using the wand tool. A [protein of interest:GAPDH] ratio was calculated and experimental conditions were normalized to mock.

### Murine strains

C57BL/6J (strain #000664, WT) and B.6129S6-Eif2ak4^tm1.2Dron^/J (strain # 008240, *Gcn2^-/-^*) mice purchased from Jackson Laboratory. Mice with mutations in both *Pkr* and *Gcn2* (“double knockout mice”) were produced as follows. B6/J;C57BL/6NCrl-Eif2ak2^em1(IMPC)Mbp/Mmucd^ mice (PKR-TKO mice) (93) were crossed to B6.129S6-Eif2ak4^tm1.2Dron^/J. The genotypes of parental mice during the breeding process were determined from tail DNAs (Transnetyx). The resulting double knockout mice are designated PKR-TKOxGcn2-/- (*Pkr^-/-^Gcn2^-/-^*). Their genotypes were confirmed by analysis of tail DNA by Transnetyx. Immunoblotting of bone marrow-derived macrophage cells derived from the PKR-TKOxGCN2-/- mice confirmed that they did not express detectable PKR or GCN2 (data not shown). Genotyping PCR was performed using the primers for *Gcn2* and *Pkr* found in **Table 3**.

**Table 3:**
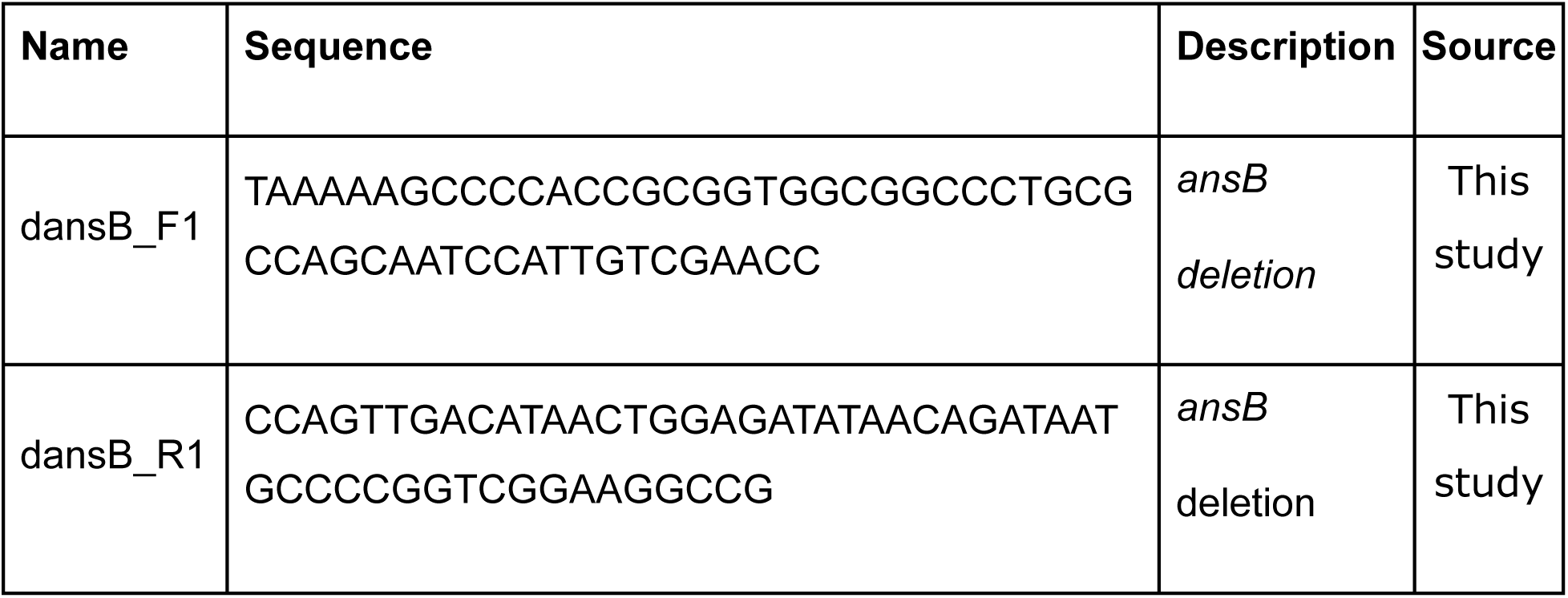
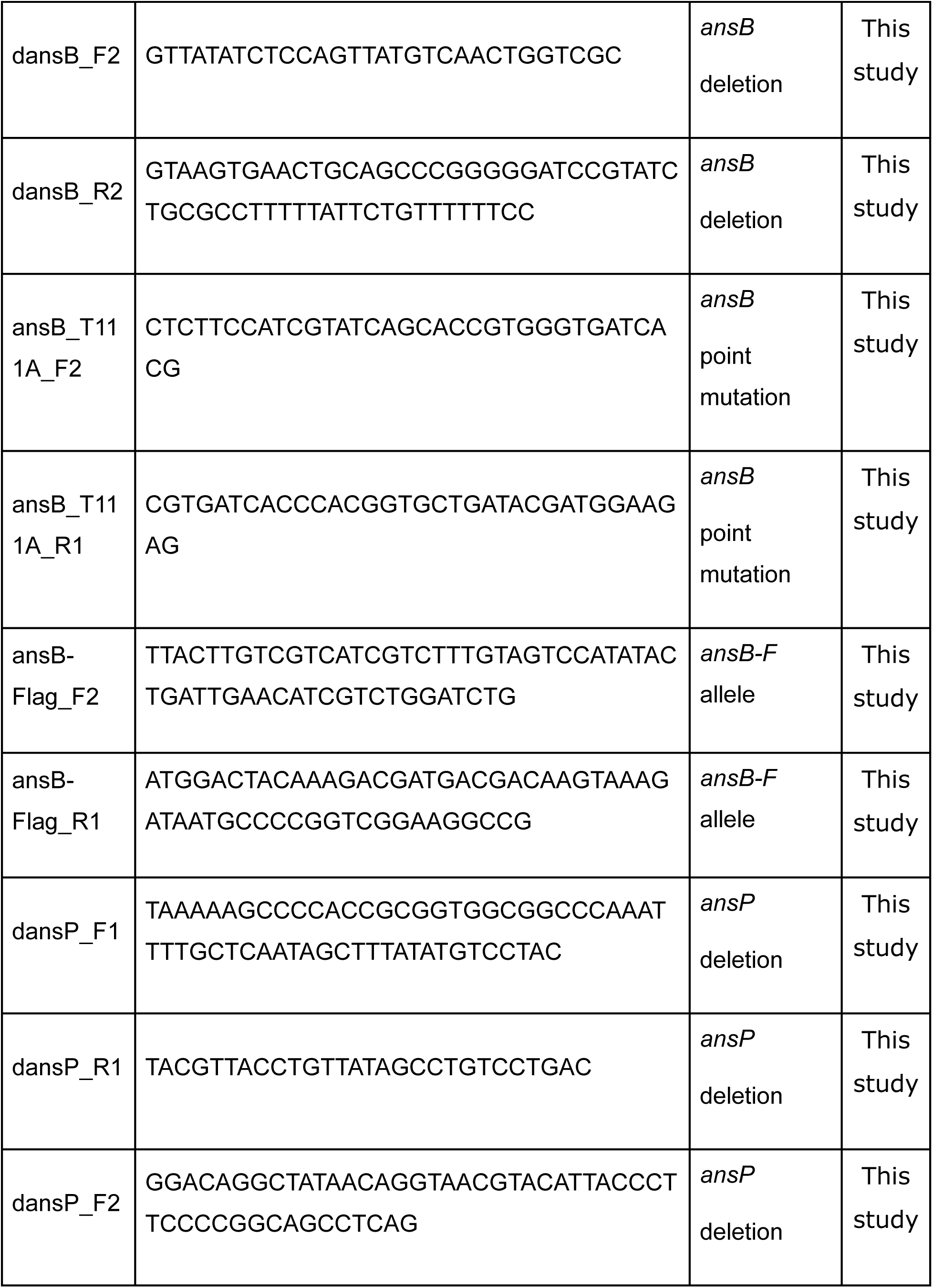
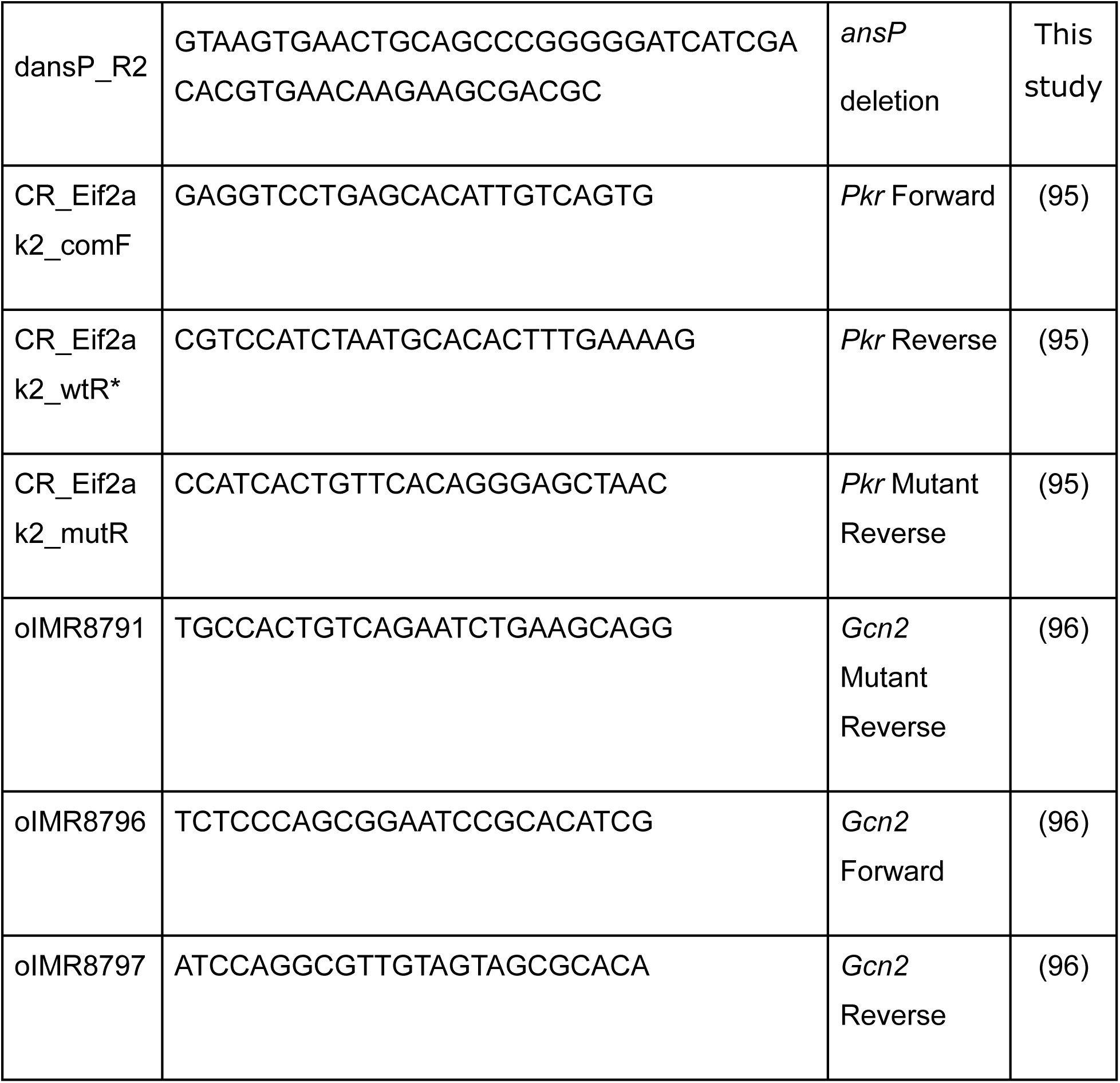
Oligonucleotides.

**Table 4:**
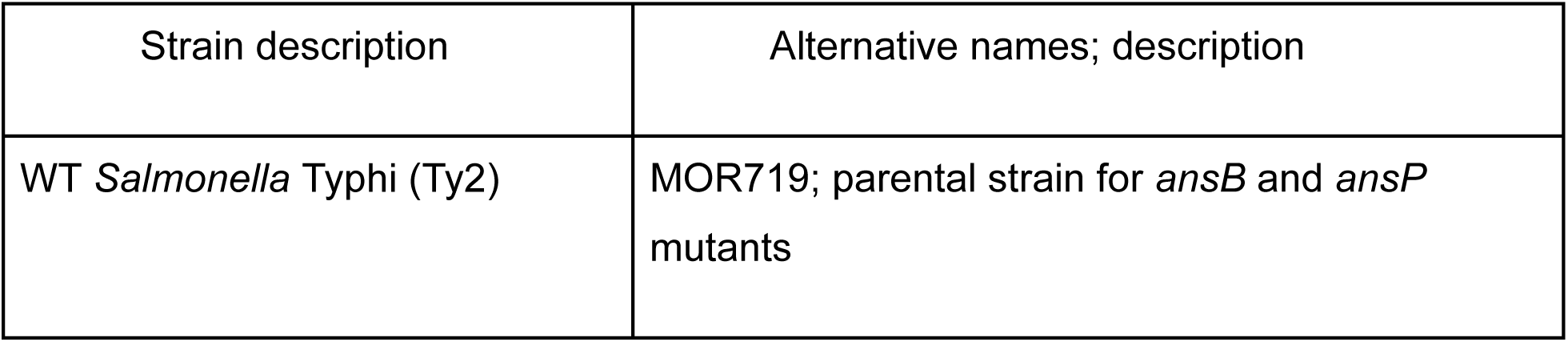
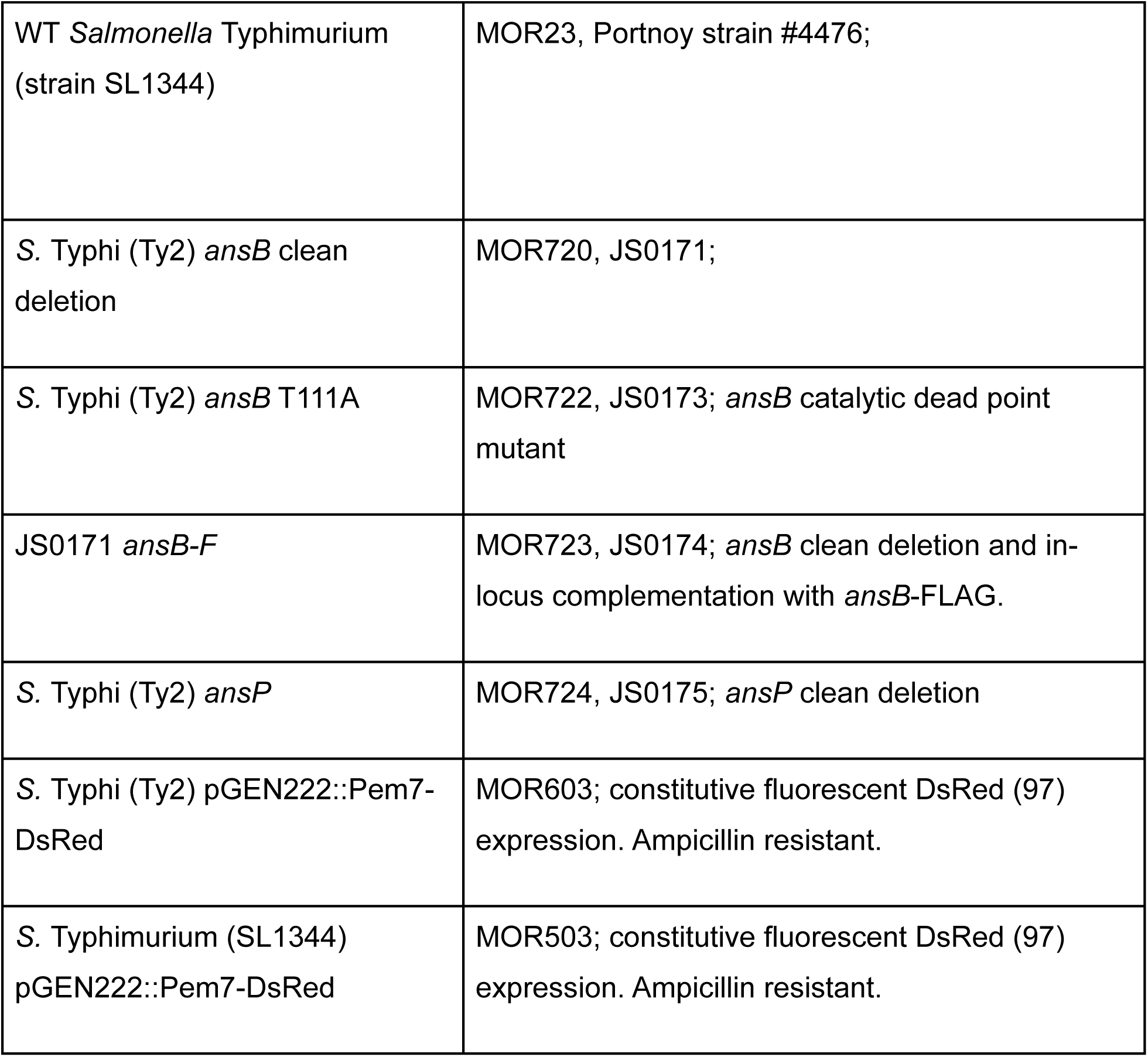
Bacterial Strains.

### Murine infections

Bacteria were cultured overnight at 37°C shaking in 3mL LB with 10mM NaCl to induce capsule expression. The following day, 1mL of pelleted culture was washed twice with PBS+/+ and diluted to the calculated density as measured by OD600 spectrophotometry. Mice were injected intraperitoneally with *S.* Typhi in 200µL PBS+/+. Inoculums are as follows: Figure 4E-G 5x10^6^ CFU/mouse; Supplemental Figure S1A-K 4x10^7^ CFU/mouse. After 24h of infection, mice were sacrificed, blood was collected in lithium heparin cuvettes via cardiac puncture and plasma isolated for cytokine analysis after separation by centrifugation at 8000rcf for 5 minutes. Organs were homogenized in tissue disruption tubes with 1.0 mm zirconia/silica beads using the Omni Bead Ruptor 24 (S=6.00; C=01; T=0:00; D=0:00) and plated on LB-agar. CFUs were counted following overnight incubation at 37°C. Outliers were analyzed by Robust regression and Outlier removal (ROUT) analysis was performed with Q=1% and excluded.

### Statistical analysis and graphing

Statistical analysis calculations were performed in Graphpad Prism v10.4.1 (627) for Windows, GraphPad Software, Boston, Massachusetts USA, www.graphpad. Where used, compact letter display comparisons note statistically significant differences p<0.05. Compact letter display statistical representation signifies differences by assigning letters between statistically different groups(94). If a letter is shared by two or more groups, the comparison is not statistically significant. If groups have no shared letters, then the comparison is significant to the tested p-value (<0.05).

### Data availability

The U-937 cell line used in this study was acquired from ATCC (catalog number CRL-1593.2) circa 2007 and validated using the ATCC Cell Line Authentication Service immediately prior to the submission of this study. The data sets presented and bacterial strains used in this study are available from the corresponding author upon request. Code generated for and used within this study for image analysis and figure preparation is made available in GitHub: https://github.com/zmpowers/software-for-GCN2.

## ETHICS APPROVAL

Murine *Salmonella* infections were performed in accordance with Protocol # PRO00012161, as approved by the University of Michigan Medical School Institutional Animal Care and Use Committee.

